# Mice lacking dopamine production in neurotensin receptor 1 neurons voluntarily undergo time-restricted feeding of high fat diet and resist obesity

**DOI:** 10.1101/2022.06.27.497778

**Authors:** Firozeh Farahmand, Michael Sidikpramana, Bara Yousef, Sarah Sharif, Kieana Shao, Qijun Tang, Gina M. Leinninger, Ali D. Güler, Andrew D. Steele

## Abstract

The introduction of processed foods high in fat and sugars has caused a dramatic increase in obesity in humans. Diet-induced obesity (DIO) can be modeled in laboratory mice by increasing the fat content of their diet. Previously, it was determined that mice lacking dopamine receptor 1 (Drd1) are completely resistant to DIO and do not eat as much food during the day as control mice. Surprisingly, when Drd1 is restored to the suprachiasmatic nucleus (SCN), which is the central regulator of circadian rhythms, these mice increase day-eating and become obese. The source of dopamine in the SCN is the ventral tegmental area (VTA), but the genetic identity of the dopamine neurons is unknown. Here we create conditional deletion mutants for *tyrosine hydroxylase* (*TH*) using *neurotensin receptor 1* (*Ntsr1*) *Cre* and other *Cre* drivers and measure feeding and body weight homeostasis on standard and high fat diets. Control mice were susceptible to DIO and overate during the day whereas *Ntsr1-Cre* conditional knockouts for TH mice did not increase day-eating, nor did they gain much weight on HFD. We used an adeno-associated virus to selectively restore TH to the VTA Ntsr1 neurons and observed an increase in body weight and increased day-eating of HFD. These results implicate VTA Ntsr1 dopamine neurons as promoting out-of-phase feeding behavior on a high fat diet that could be an important contributor to diet-induced obesity.

## Introduction

Circadian (∼24h) rhythms of physiology and behavior are regulated at the cellular level by transcriptional and post-translational feedback loops of core clock proteins and regulatory RNA [1]. This process is cell-autonomous and maintains its period in the absence of external time cues. In mammals, a population of neurons in the suprachiasmatic nucleus (SCN) of the hypothalamus serves as the “master clock”, which synchronizes the autonomous clocks of both central and peripheral tissues [2]. Light and other external factors (e.g. food availability) modulate the expression of clock genes to reset and entrain the timing of the circadian clock in different tissues [1]. Mounting evidence demonstrates that circadian rhythms are an integral part of the behavior and physiology related to energy homeostasis [3, 4].

The relationship between metabolism, reward circuitry, and circadian rhythmicity is poorly understood [5]. A growing body of evidence suggests that eating out-of-phase from light entrained circadian rhythms is a driver of obesity [6, 7]. We recently demonstrated that dopamine receptor 1 (Drd1) signaling promotes out-of-phase feeding on a high fat diet (HFD) and leads to diet-induced obesity (DIO) in mice [8]. Drd1 global knockout (Drd1-KO) mice do not increase day-eating of HFD, resist DIO, and do not show inflammatory phenotypes associated with DIO. Remarkably, the rescue of Drd1 expression solely in the SCN of Drd1 KO mice restored susceptibility to day-eating of HFD, DIO, and other associated metabolic disorders [8].

Ventral tegmental area (VTA) dopamine neurons send functional projections to the SCN [9]. For example, selective activation of dopamine neurons in the VTA increases the rate of photoentrainment to shifting light schedules [9]. The neuropeptide neurotensin (Nts) is important in the regulation of various physiological functions including sleeping, eating, and drinking behaviors [10, 11]. Nts has two different receptors, termed neurotensin receptors 1 and 2 (Ntsr1 and Ntsr2)--with Ntsr1 having a prominent role in the regulation of body weight and feeding. Interestingly, one of the major sites of Ntsr1 expression is the VTA. Here we used conditional deletion of tyrosine hydroxylase (TH), the rate limiting enzyme in DA production [12], in Ntsr1 expressing and other dopamine populations to probe the function of various dopaminergic neurons and the requirement of dopamine release for promoting day-eating associated with DIO. Our data presented here suggest that the Ntsr1 dopamine neurons of the VTA promote day-eating and DIO associated with rewarding, energy-dense foods.

## Results

To identify the neurons releasing dopamine into the SCN to drive out-of-phase palatable food consumption and DIO, we used a panel of *Cre*-driver lines to disrupt a conditional allele of *TH* [13]. Genetic ablation of the *TH* gene prevents dopamine production in targeted neurons and hence eliminates their dopaminergic identity while leaving their other neurotransmitter and neuropeptide systems intact, providing a strategy to probe the necessity of dopamine in a given circuit [14]. In preliminary studies, conditional knockout (cKO) lines using either *Ntsr1-Cre* [15], *Slc17a6* (*vesicular glutamate transporter; Vglut2*)*-Cre* [16], *Calbindin1 (Calb)-Cre* [17], *Cholecystokinin*(*CCK*)-*Cre* [18], or *Brain-derived neurotrophic factor* (*BDNF*)*-Cre* [19] were tested for weight gain on standard chow diet (SCD) and HFD. We observed that most cKO lines gained a similar amount of weight upon being transitioned to a HFD for 6 weeks as did pooled controls (**Fig. 1 and Supp. Fig. 1**). However, the *Ntsr1-Cre* cKO showed marked resistance to DIO; 2-way ANOVA revealed significant effects of time (p<0.0001), genotype (p=0.0036), and time x genotype (p<0.0001). The *Ntsr1-Cre* cKO mice had significantly lower body weight at weeks 4-8 of the study (week 4 p=0.0018, week 5-8 p<0.0001) and *Vglut2-cre* cKO also showed reduced weight gain (Bonferroni’s multiple comparison tests p=0.0027 for week 5, p=0.0007 for week 6, P=0.0005 for week 7, and p=0.0002 for week 8). Notably, when *Vglut2-Cre* cKO mice were compared against littermate control mice rather than pooled controls, there were no significant difference in normalized body weights on HFD (p>0.999) (**Supp. Fig. 1**). *BDNF-Cre* cKO showed a small but significant decrease in normalized body weight compared to littermate controls at weeks 6-8 of the study (p=0.013 for week 6, p=0.0017 for week 7, and p=0.0013 for week 8, 2-way ANOVA with Bonferroni multiple comparisons) (**Supp. Fig.1**).

**Figure 1.**
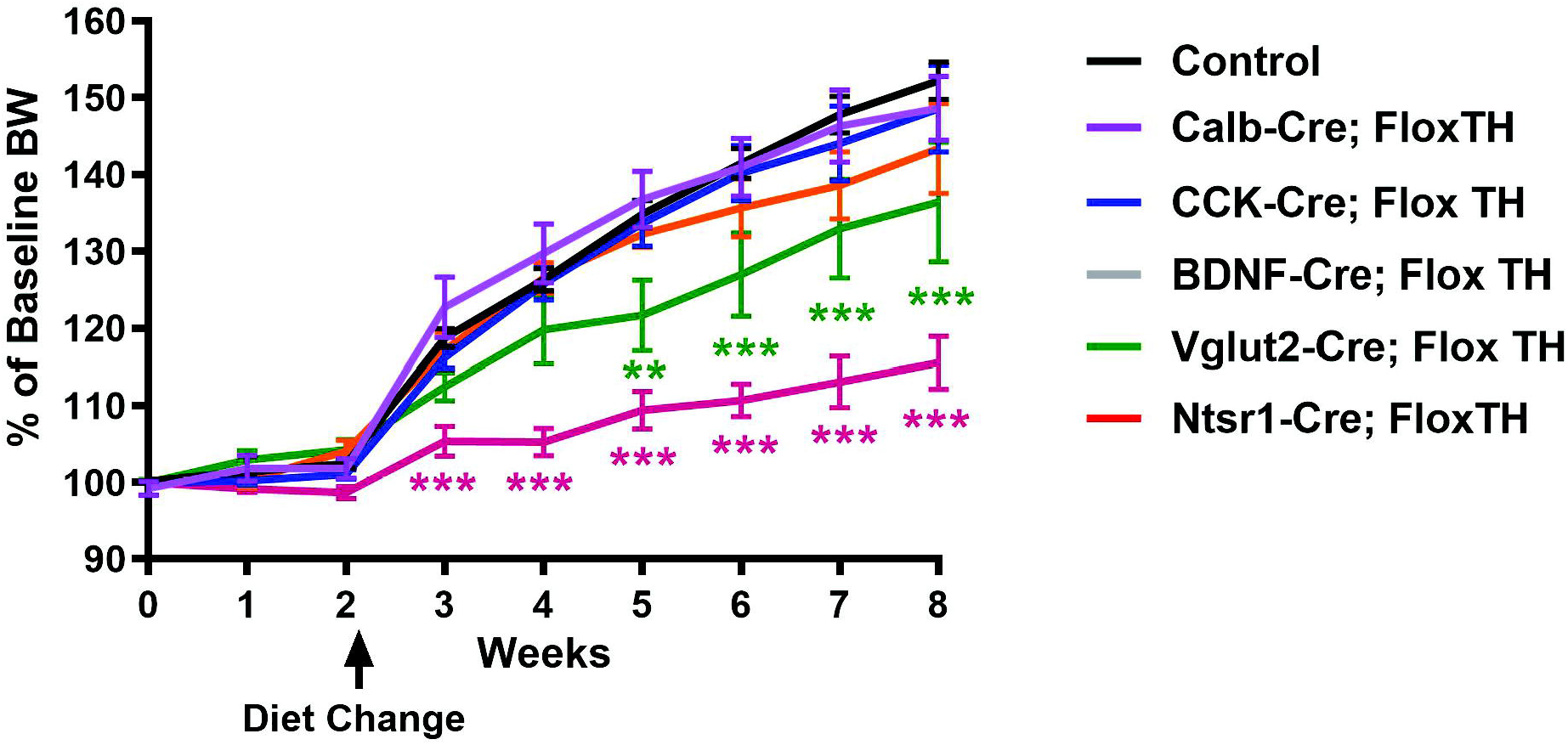
Screen for dopamine mutants with altered diet-induced obesity. Body weight measurements of pooled controls (n=37), BDNF-Cre (n=6), Calb-Cre (n=6), CCK-Cre (n=5), Ntsr1 (n=8) and Vglut2-Cre (n=7). The first three measurements were made on SCD and afterwards the mice were transitioned for HFD for the next 6 weekly measurements. (2-way ANOVA with Bonferroni **p<0.01, ***p<0.001)

For further characterization, we chose to study the *Ntsr1-Cre* cKO line given that deletion of *TH* using *Ntsr1-Cre* markedly impairs weight gain on an HFD without affecting weight on SCD (**Fig. 2A and Supp. Fig. 2**). The percent change in BW for male control mice was about 45% of initial weight whereas *Ntsr1-Cre* cKO mice only increased weight by 15% after 6 weeks of HFD (**Fig. 2A**) (2-way ANOVA mixed-effects analysis genotype comparison p-value = 0.0005 [F{1, 8}=31.29]; Bonferroni multiple comparisons, weeks 0-3 p>0.9999, week 4 p=0.0018, weeks 5-8 p<0.0001). In a pilot study of female *Ntsr1-Cre* cKO mice, similar results were obtained on HFD (**Supp. Fig. 2C-D**). In terms of food intake, there were several significant fixed effects (e.g., night vs. day p<0.0001, SCD vs HFD p =0.0026 whereas control versus cKO p=0.237). On HFD, control mice significantly increased their day-eating in comparison to SCD (p=0.0011, mixed-effect analysis, Bonferroni correction); however, *Ntsr1-Cre* cKO mice did not increase day-eating of HFD (P>0.999, mixed effect analysis, Bonferroni correction) (**Fig. 2B**). Neither control (p=0.0519) nor *Ntsr1-Cre* cKO (p>0.999) mice showed increased night food intake when switched from a SCD to HFD (**Fig. 2B**). The resistance of *Ntsr1-Cre* cKO to DIO cannot be explained by hyperactivity because testing of home-cage behavior using video-based computer vision software [20] showed that their high activity behaviors (jumping, hanging, walking and rearing) were not increased on HFD (one-way ANOVA p=0.0159, multiple comparisons test with Bonferroni correction, control SCD versus control HFD p=0.354; Ntsr1-Cre cKO SCD vs HFD p=0.849) (**Fig. 2C**). We visualized the deletion of TH within the VTA of the *Ntsr1-Cre* cKO mice by staining for DAT and TH, determining that this line deletes TH in ∼70% of dopamine neurons in VTA: on average, control mice had 184 TH+ cells per section while the *Ntsr1-Cre* cKO mice had 53 (p-value of p<0.0001, unpaired t-test) (**Fig. 2D**). To test for off-target deletion of TH in other catecholamine nuclei important for regulating feeding and energy homeostasis [21], we examined dopamine populations in the zona incerta, arcuate nucleus, dorsomedial hypothalamus, and locus coeruleus, finding them to be largely intact, consistent with prior detailed characterizations of the *Ntsr1-Cre* line (**Supp Fig. 2B**) [15, 22].

**Figure 2.**
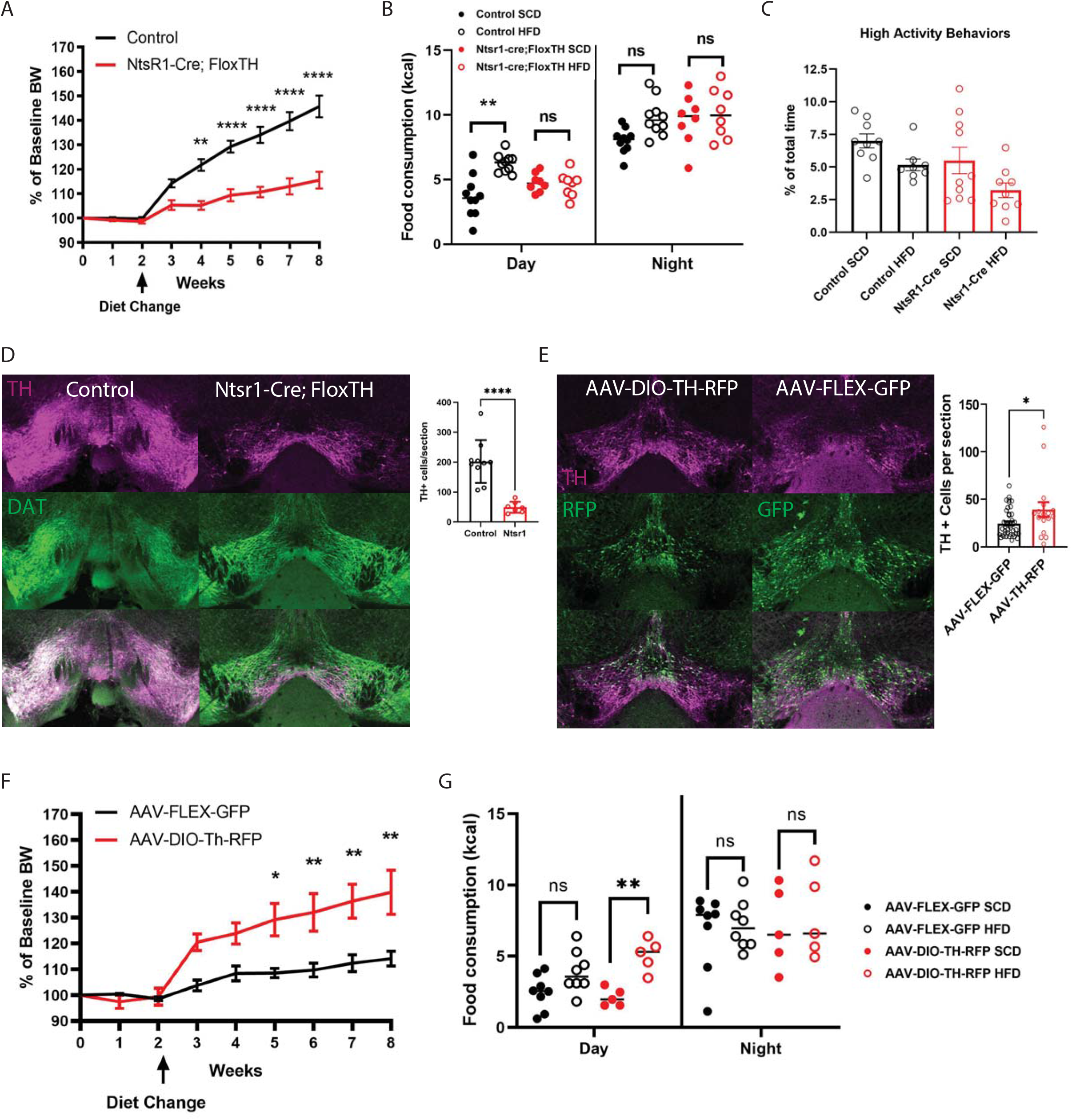
*Ntsr1-Cre* deletion of TH. (A) Body weight measurements of controls (n=10) and *Ntsr1-Cre* (n=8). (B) Day-versus night-eating in *Ntsr1-Cre* cKO mice on SCD and HFD (n=8-9 per group; 3-way ANOVA with Bonferroni’s multiple comparisons test **p<0.01). (C) High activity behaviors in the home-cage (jumping, rearing, hanging, and walking) on SCD and HFD. (D) TH and DAT staining in the VTA of *Ntsr1-Cre* cKO and control VTA and cell counts (****p<0.0001, unpaired t test). (E) TH and RFP staining in the VTA of an *Ntsr1-Cre* cKO mouse injected with AAV-DIO-TH-RFP, TH and GFP staining in the VTA of an *Ntsr1-Cre* cKO mouse injected with AAV-FLEX-GFP, and the average number of TH positive cells in AAV-FLEX-GFP and AAV-DIO-TH-RFP injected *Ntsr1-Cre* cKO mice (p<0.05, unpaired t test). (F) Body weight measurements of AAV-FLEX-GFP (n=8) and AAV-DIO-TH-RFP (n=5) injected *Ntsr1-Cre* cKO mice on SCD and HFD. (G) Day-and night-eating in AAV-FLEX-GFP and AAV-DIO-TH-RFP injected *Ntsr1-Cre* mice (n=5-8 per group; 3-way ANOVA with Bonferroni’s multiple comparisons test, **p<0.01).

To solidify that it was the VTA Ntsr1 dopamine neurons, as opposed to other sites, mediating the effects on day-eating and DIO, we utilized an AAV Double-Floxed Inverted Open (DIO) reading frame encoding TH-RFP to selectively restore TH to the VTA of *Ntsr1-Cre* cKO adult mice. Control *Ntsr1-Cre* cKO mice were given AAV DIO-GFP injections into the VTA. To verify the restoration of TH by AAV-DIO-Th-RFP within the VTA of the *Ntsr1-Cre* cKO mice, the number of TH+ cells and fluorescent+ cells (either GFP or RFP depending on which virus was administered) is shown in both a AAV-Flex-GFP and AAV-DIO-Th-RFP injected mice (**Fig. 2E**). An overlap of TH and the fluorescence is also shown to demonstrate the increase in TH found in AAV-DIO-Th-RFP injected mice in comparison to the AAV-Flex-GFP mice (**Fig**. is a result of the AAV-DIO-Th-RFP injection (**Fig. 2E**). To compare the staining, the average number of TH+ cells per section (n=4-8) sections per mouse from a rostral to caudal sample) of VTA was analyzed (**Fig. 2E**). On average, the AAV-Flex-GFP injected mice had 25 TH+ cells per section while the AAV-DIO-Th-RFP mice had 39 (p<0.0001, unpaired t-test). The percent change in BW showed that mice injected in the VTA with AAV DIO-Th-RFP increased their BW substantially, gaining about 30% of initial weight whereas AAV-FLEX-GFP injected mice only increased weight by about 15% (**Fig. 2F**). The effect of genotype was strong (p=0.0066 (F(1, 7)=14.56); there was a difference between percent of baseline body weight for weeks 5-8 (week 5 p=0.0275 week 6 p=0.0099, week 7 p=0.0035, week 8 p=0.0013 Bonferroni multiple comparisons). As expected, AAV-DIO-TH-RFP injected mice increase their daytime eating when switched to HFD (p=0.0014, Bonferroni multiple comparisons), whereas the *Ntsr1-Cre* cKO mice injected with control AAV-Flex-GFP did not increase day eating on HFD (p>0.9999; **Fig. 2G**). There were no significant differences between night food intake on SCD and HFD, as expected (AAV-DIO-Th-RFP p>0.9999, AAV-Flex-GFP p>0.9999).

## Discussion

In summary, our genetic strategy allows for identification of dopamine subsets required for day-eating and DIO. Our analysis demonstrated that dopamine production in Ntsr1 dopamine is needed for DIO. When placed on a HFD, Ntsr1-Cre cKO adult mice do not become obese or experience an increase in calorie intake during the day in contrast to their wild-type littermate controls. We determined that VTA Ntsr1 dopamine neurons mediate these effects by using AAV to selectively restore TH to these neurons in an otherwise cKO background, finding that cKO mice with restored expression of TH in the VTA gained more weight and were subject to increased day-eating in comparison to controls on a HFD. How do energy-dense diets disrupt the circadian regulation of meal timing to exacerbate their deleterious effects on metabolism and body weight regulation? We hypothesize: (1) rewarding foods increase out-of-phase food consumption by perturbing central circadian control, (2) they accomplish this by engaging the dopaminergic VTA→SCN circuit, and (3) the dopamine-dependent SCN signaling regulates hedonic meal timing. Increased dopaminergic tone on the circadian pacemaker dampens SCN neuronal activity (by GABAergic SCN neurons), disinhibiting downstream orexigenic targets, and ultimately promotes out-of-phase foraging [23, 24]. Several hypothalamic nuclei are downstream targets of the SCN; the dorsomedial hypothalamus (DMH), lateral hypothalamus (LH) and paraventricular nucleus (PVH), have been implicated in the circadian regulation of feeding. The DMH, a nucleus that limits feeding by inhibiting arcuate AgRP neurons, receives direct and indirect inputs from the SCN and its lesion abolishes food intake rhythms [25]. The SCN via indirect inputs to the LH drives circadian activity and orexin release associated with arousal and feeding. The LH lies at the interface of feeding and reward processing [26]. The SCN also signals to the PVH, the primary target of arcuate AgRP neurons, via diffusible factors (i.e. arginine vasopressin) and direct projections, to entrain the circadian oscillations of various metabolically relevant hormones (e.g. corticotropin-releasing hormone and oxytocin) [27]. Notably, the Arc contains a circadian clock and makes reciprocal connections with the SCN and its targets to ensure coordinated daily metabolic synchrony [28]. Although a wealth of communication between the SCN and centers that control food intake and metabolism has been discovered, the circuits that control meal timing and the underlying neurophysiological mechanisms have not been characterized.

Our findings define a novel component of the feeding circuit and will provide therapeutic targets in the fight against obesity. Animals on a HFD exhibit a profound disruption in the circadian regulation of food intake; they increase rest phase food consumption without reducing active phase consumption. Overconsumption of calories under reduced RER together with dampened circadian rhythmicity favors increased fat storage and weight gain [29]. This behavior is akin to humans snacking on junk food outside of mealtimes resulting in dampened fat utilization between meals [7]. Our data suggest that Ntsr1 marks the dopamine neurons that are required for DIO; so it stands to reason that the Ntsr1-dopamine neurons project to the SCN to mediate day eating, but SCN specific manipulations of Ntsr1 dopamine have yet to be conducted. Although our lead candidate for the source of SCN-dopamine in response to HFD is VTA-neurons, which play a central role in reward processing, it remains possible that other nuclei also provide dopamine to the SCN in response to palatable foods. Future studies will use viral tracing methods to characterize the projections of Ntsr1 VTA SCN neurons, manipulate these neurons using opto-or chemogenetics, and use Cre driver lines with more restricted expression to the VTA→SCN circuit to test the effects on feeding and body weight homeostasis.

The advent of targeted meal timing to treat obesity is elegant in its simplicity. While early findings are promising, the field lacks a mechanistic explanation for why such a treatment might work. Moreover, it is becoming clear that unhealthy eating habits exacerbate metabolic disorders by disrupting circadian rhythms. Continued investment in research on the circuitry and neurotransmitter systems required for this pathological feedforward loop of dopamine to SCN to metabolic disorder. Such studies will provide a more refined understanding of the relationship between hedonic feeding and circadian circuits, and provide an important basis for future chronotherapeutic interventions.

## Methods

The mouse lines used in this study were obtained from Jackson labs or from donating investigators; we obtained *Vglut2-Cre* (016963), *CCK-Cre* (012706), *BDNF-Cre* (030189), *Calb-Cre* (028532) from Jackson labs; *Ntsr1-Cre* (available at Jackson labs as stock 033365) mice came from the breeding colony of GL, and TH flox/delta mice from Richard Palmiter (University of Washington) [13]. Tail clips were sent to Transnetyx® (Cordova, Tennessee) for qPCR analysis of the following alleles: *Cre, TH* delta allele, *TH* floxed allele, and *TH* wild-type allele. Male mice were used for most experiments (except Fig S2C-D). Mice were weaned onto SCD (18% fat, 24% protein, and 58% carbs; Envigo, 2018). At 9-10 weeks of age, mice were singly housed in enriched small static microisolator cages, including a cotton nestlet, Bed-r’Nest (Lab Supply), and a Safe Harbor Mouse Retreat (Bio-Serv). After single housing BW, was measured weekly for three weeks on SCD, followed by six weeks on HFD (59% fat, 15% protein, and 26% carbohydrates, BioServ). Measurement of day versus night eating was conducted on both an SCD (∼10 days after single housing) and an HFD (∼10 days after transitioning diet). For these measurements, three days and three nights of food intake values were recorded and averaged. To compute calorie intake for SCD, the grams of food recorded were converted to 3.1 kcal/g whereas the value of 5.49 kcal/g was used for HFD.

Stereotaxic delivery of AAVs was used to restore TH in a Cre-dependent manner to restore expression of TH in *Ntsr1-Cre* cKO mice. AAV-DIO-TH-RFP was injected to restore TH and AAV-FLEX-GFP was injected as a negative control. Mice undergo surgery at 8-12 weeks of age. A rostral to caudal skin incision is made with a sterile scalpel to expose the skull. Then, the skull is leveled (Kopf, Model 1905 Stereotaxic Alignment Indicator) and bregma is identified. Two small craniotomies of 0.5mm in diameter above each brain hemisphere are made with a miniature drill at stereotaxic coordinates for the VTA (relative to bregma): x: +/-0.5mm, y: −3.7mm, z: −4.0mm. Sterile glass pipettes are inserted through the holes in the skull and 0.75μl of AAV was delivered at 10 μl/second. Once both injections are complete, bone wax (Surgical Specialties, #309) is applied to the craniotomy sites on the skull. 0.1 ml (1mg/kg of 0.25%) of Bupivacaine is given subcutanesouly and Marcaine (NDC 0409-1610-50) solution is applied on the wound margin prior to closure. The skin is closed with 0.1 ml tissue adhesive (Bausch + Lomb, 24208-780-55). 0.5 ml sterile normal saline is delivered subcutaneously before and after the procedure to ensure adequate hydration.

Immunofluorescence antibody staining was performed using chicken polyclonal TH antibody (Aves, TYH), rat monoclonal DAT (Invitrogen, MAB369), and anti-RFP (Takara, 632543) primary antibodies. An Alexa Fluor 647-conjugated Affinipure Goat Anti-Chicken IgG (Invitrogen) and an AffiniPure Goat Anti-Rat 568 IgG (Invitrogen) or Goat Anti-Rat 488 IgG (Invitrogen) were used as secondary antibody staining. Imaging was conducted via a Nikon A1 confocal microscope linked to a computer with NIS-Elements BR 3.0 software, and image analysis and enhancement was performed with ImageJ (Fiji) software on a separate computer system. To quantify neuron, we sectioned n=4=6 brains per group and sampled n=4-8 sections per mouse from a rostral to caudal sampling of midbrain.

## Figure legends

Figure S1. Average weight in grams and normalized body weights for the lines examined.

Figure S2. (A) Weight in grams for Ntsr1-Cre cKO mice. (B) TH staining in zona incerta, arcuate nucleus, dorsomedial hypothalamus, and the locus coeruleus. (C) Body weight for female (n=4 WT and n=5 cKO) mice on SCD and HFD in grams. (D) Normalized body weight for the same mice in (C).

