## Supplementary figures and images for "Mice lacking dopamine production in neurotensin receptor 1 neurons voluntarily undergo time-restricted feeding of high fat diet and resist obesity"

### Supplemental Figure 1

Supplemental Figure 1

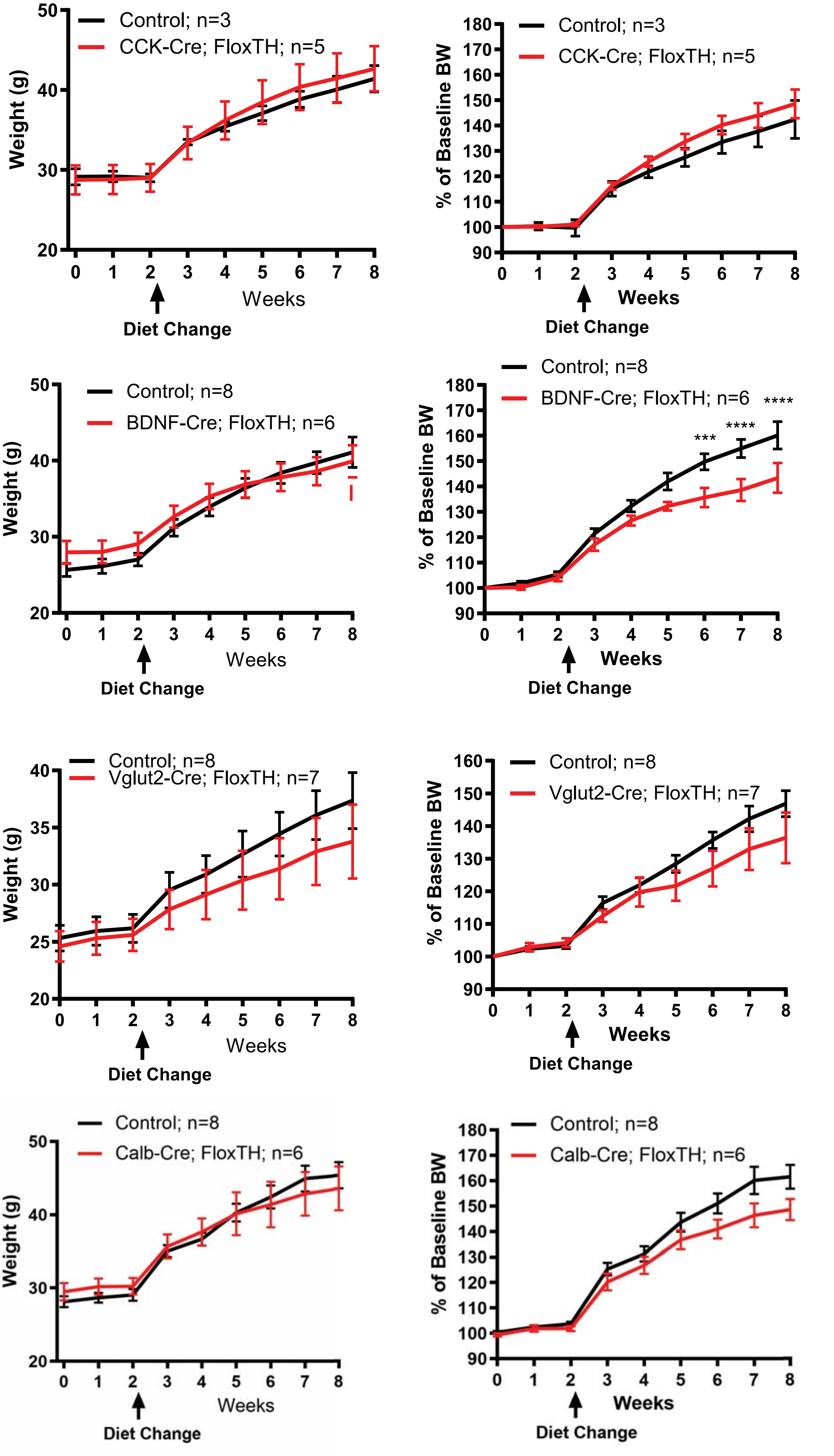

### Supplemental Figure 2

Supplemental Figure 2

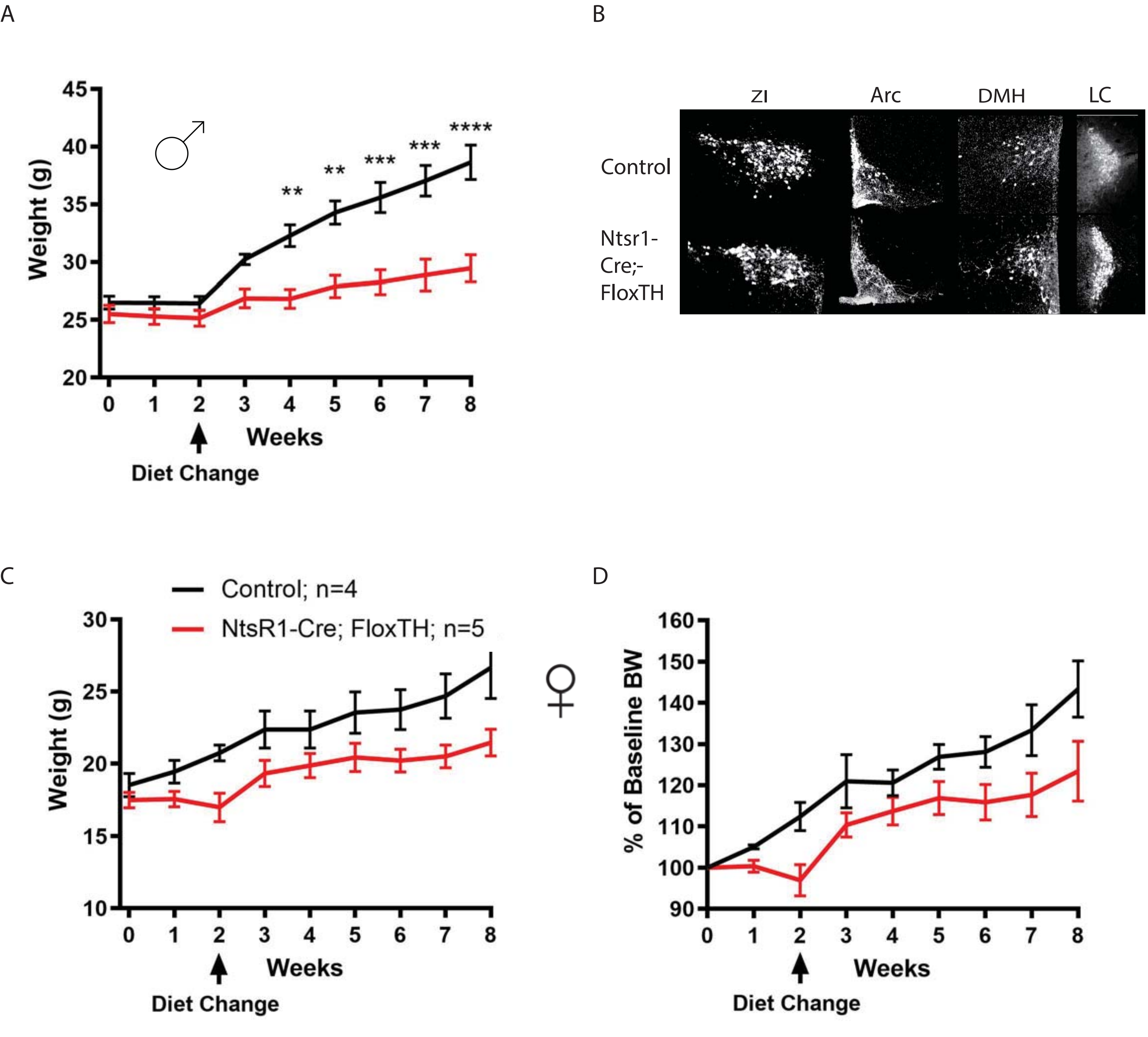
